# 3D-printed decoys are as effective as taxidermied decoys in attracting red-breasted sapsuckers for mist-netting

**DOI:** 10.1101/2022.03.27.485678

**Authors:** Libby Natola, Frankie Tousley

**Affiliations:** University of British Columbia, 2329 West Mall, Vancouver, BC V6T 1Z4, Canada; Indiana State University, 200 N 7th St, Terre Haute, IN 47809, United States

**Keywords:** 3D-printed decoys, taxidermied decoys, mist-netting, trapping, field ornithology

## Abstract

Decoys often improve targeted mist-netting efforts by drawing the species of interest to a specific area nearer the net. Traditional decoy constructions include taxidermied carcasses, hand-made wood or clay figures, or professionally made products purchased from companies that provide a limited number of species, sizes, shapes, and markings. 3D-printing allows ornithologists to customize decoys to their own study species’ specifications using cheap, durable, and replaceable materials. We show that red-breasted sapsuckers (*Sphyrapicus ruber ruber*) respond with equivocal aggression towards 3D-printed decoys and taxidermied decoys, demonstrating 3D-printed decoys as an effective tool in attracting birds towards a mist net for capture.

## Introduction

Many facets of ornithological research require the physical capture of birds. Even with targeted mist-netting, it can be challenging to capture a particular species of interest. In effort to improve capture rates, decoys made from taxidermied conspecifics are deployed near mist nets to lure territorial birds closer (Angelier et al. 2010; Slevin et al. 2016). Despite their efficacy, good decoys can be difficult to source. Carved wood, stuffed fabric, papier mache, and sculpted clay decoys (Slevin et al. 2016; Heward et al. 2017; de Zwaan et al. 2021) are sometimes used, but require artistic skill to render a lifelike replica capable of attracting a bird. This may be beyond the talents of many biologists. These decoys are often fragile, susceptible to water damage, and may pose a risk of injury to particularly aggressive birds (Slevin et al. 2016). Some decoys are available for purchase, but these are often limited in species. Decoys may also be made from taxidermied skins of the study species (Angelier et al. 2010; Seneviratne et al. 2012; Bentz et al. 2019), but these are fragile and may be dependent on access to a museum, collection permits, or require euthanizing a bird. This option may be particularly out of reach for researchers of understudied or at-risk species.

3D-printing is increasingly accessible and presents an alternative that overcomes many of the aforementioned obstacles associated with traditionally used decoys. 3D-printed decoys can be manufactured without any specialized artistic skills, may be customized to depict any species, and do not require a dead specimen; they are also affordable, durable, weather resistant, and readily replaceable. Further, their design can facilitate moving mechanical components that mimic movement or aggressive postures which may attract or provoke birds. 3D-printed decoys have successfully facilitated research on beetles (Domingue et al. 2014) and turtles (Bulté et al. 2018). In avian research, 3D-printed decoys enabled egg-rejection (Igic et al. 2015) and passerine territoriality studies (Beck et al. 2019; Bentz et al. 2019), but the effectiveness of entirely plastic decoys has not been evaluated in avian mist-netting. We have designed a 3D-printed decoy and tested it against a taxidermied decoy for efficacy in attracting red-breasted sapsuckers (*Sphyrapicus ruber ruber*).

## Methods

### Taxidermied decoy production

Libby Natola made the taxidermied decoy from a red-breasted sapsucker carcass donated to the Beaty Biodiversity Museum (CWS permit BC-SA-0022-16). She skinned the carcass, stuffed and posed it around a cotton and metal wire armature, then mounted it to a small branch in a lifelike position. With the goal of improving decoy detection and allure, we attached the branch mount to a pulley system (Figure 1a) to broadcast the decoy’s location and increase decoy credibility through behavioural mimicry.

**Figure 1.**
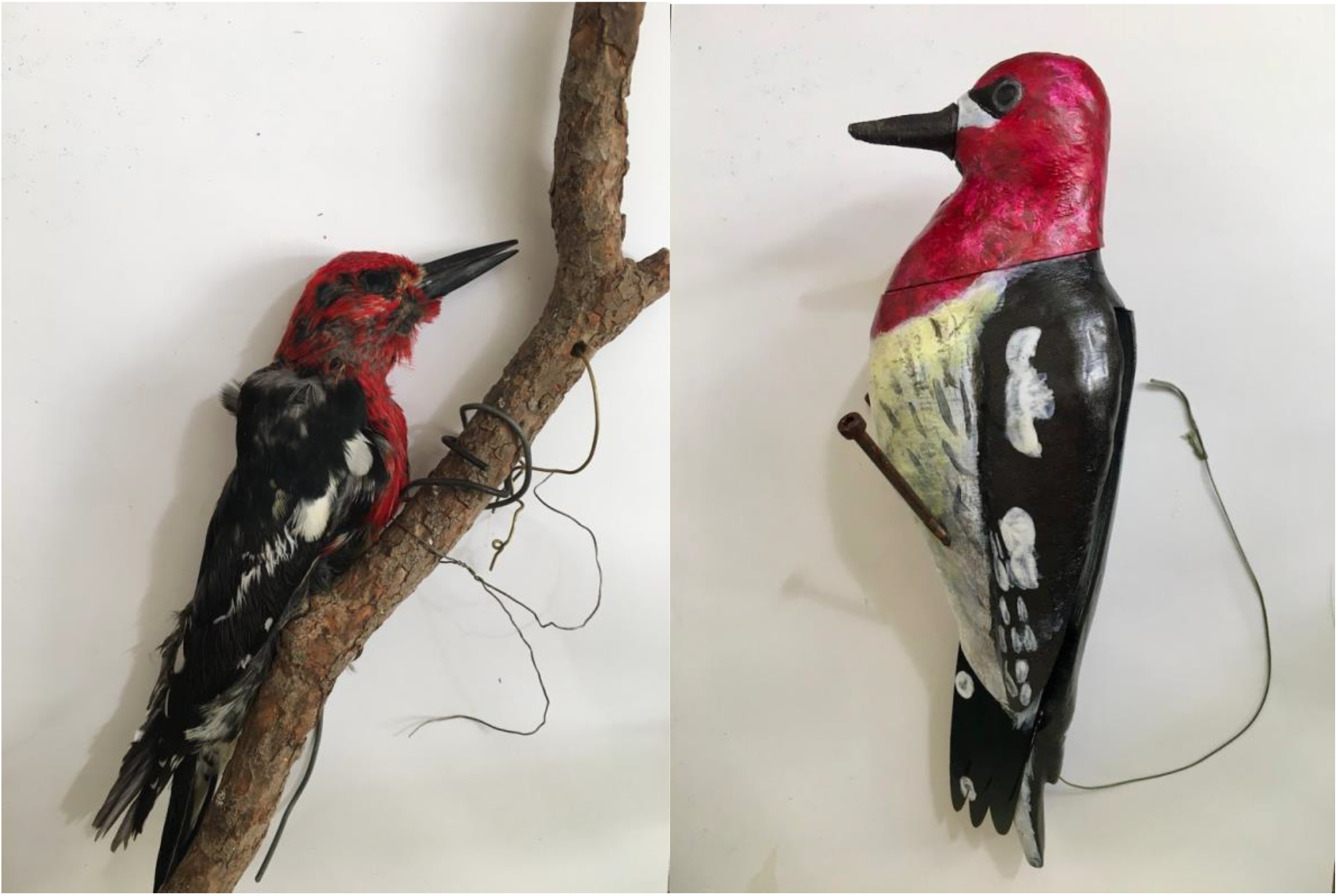
Taxidermied (a) and 3D-printed (b) decoys used in this study.

### 3D-printed decoy production

Frankie Tousley designed and manufactured the plastic decoy using 3D computer-aided design and printing software; we used only free and open-source software, or software available at no cost under personal-use licensure. We used Blender 3D creation software (version 2.8 beta, Blender Foundation 2018) to digitally sculpt the blank decoy, then Fusion 360 (Autodesk, Inc., USA) to develop the decoy’s mobile, mechanical elements. The completed decoy was prepared for printing in Slic3r Prusa Edition (version 1.4.3, 2019), and printed on a fusion deposition modeling (FDM) 3D printer in polyethylene terephthalate glycol-modified (PETG) plastic. The final design consisted of 17 printed pieces, assembled then painted with acrylics (Figure 1b). We incorporated an internal pulley system in the decoy design. Unlike the vertical movement of the taxidermied decoy, the plastic decoy pulley system unfurls folded wings into an aggressive posture common of woodpeckers. As with the taxidermied decoy, this behavioural mimicry might alert or entice a bird targeted for capture.

### Field assessment

We performed all decoy trials on red-breasted sapsuckers in the Prince George and Vancouver areas of British Columbia, Canada from May–June of 2019 and 2020. We identified sapsucker territories either by the presence of fresh sap wells, or by sapsucker reactions to audio playback of conspecific calls. Upon confirming sapsucker presence, we conducted a trial within the defended territory boundary. During each trial we hung one decoy from a tree trunk, approximately two meters high, in a location of little vegetation to maximize decoy visibility. We determined decoy selection for the first trial by coin flip, then alternated decoys for all subsequent trials. One camouflaged technician controlled the decoys’ movement as described herein via pull string. Concurrently, we ran a 10-minute standardized audio playback from a speaker placed at the base of the decoy tree; playback included a series of red-breasted sapsucker drumming bouts, aggression calls, and silences. Author LN narrated sapsucker reactions and movement into an audio recorder for later analysis. As a standard for measurement, we delineated two concentric zones around the decoy at 2 m and 10 m radii. Data collected include number of approaches by the trial sapsucker into the 2 m and 10 m zones, number of approaches to the decoy tree, nearest approach distance to the decoy (m), number of apparent attacks on the decoy, and success or failure of capture. LN estimated all distance measurements visually.

### Statistical Analysis

We conducted two-sample t-tests comparing the effect of decoy type on observations for a trial sapsucker’s nearest approach distance, and the number of times they landed within the 2 m and 10 m zones. We performed Fisher’s exact tests for the effect of decoy type on the number of trial sapsuckers that approached into the 10 m zone, as well as the number that attacked the decoy. We also performed chi-squared tests to compare the effect of decoy type on the number of trial sapsuckers that approached into the 2 m zone, as well as the number that landed on the decoy tree.

## Results

We conducted 43 total trials, though in two trials the target sapsucker exited the area after initial detection. The 41 remaining trials that we subjected to statistical analysis involved sapsuckers that responded to audio playback with aggressive territorial behaviour. We found a significant difference between the two decoy types for the nearest approach distance by sapsuckers (two-sample t-test t = 2.3, p = 0.03); the mean nearest approach distance by sapsuckers was 2.4 m for printed decoys, and 4.6 m for taxidermied decoys (Figure 2). This was the only statistically significant difference found in sapsuckers’ responses to our decoy treatment. Sapsuckers were equally likely to land on the decoy tree (X-squared test, X-square = 0.01, p = 0.91) or attack a decoy (odds ratio = 0.51, Fisher’s exact test p = 1), regardless of decoy type. Similarly, the decoy types produced no significant difference in the frequency (per trial) that sapsuckers landed on the decoy tree (two-sample t test, t = 1.47, p = 0.15) (Figure 3). There was no significant difference in the likelihood that a sapsucker would approach into the 2 m zone (X-squared test, X-squared = 0.37, p = 0.54) or 10 m zone (Fisher’s exact test, p = 0.49) of either decoy type, and the frequency (per trial) that sapsuckers approached into the 2 m zone (t-test, t = 1.42, p = 0.17) or 10 m zone (t-test, t = 1.57, p = 0.13) (Figure 3) was also equivalent.

**Figure 2.**
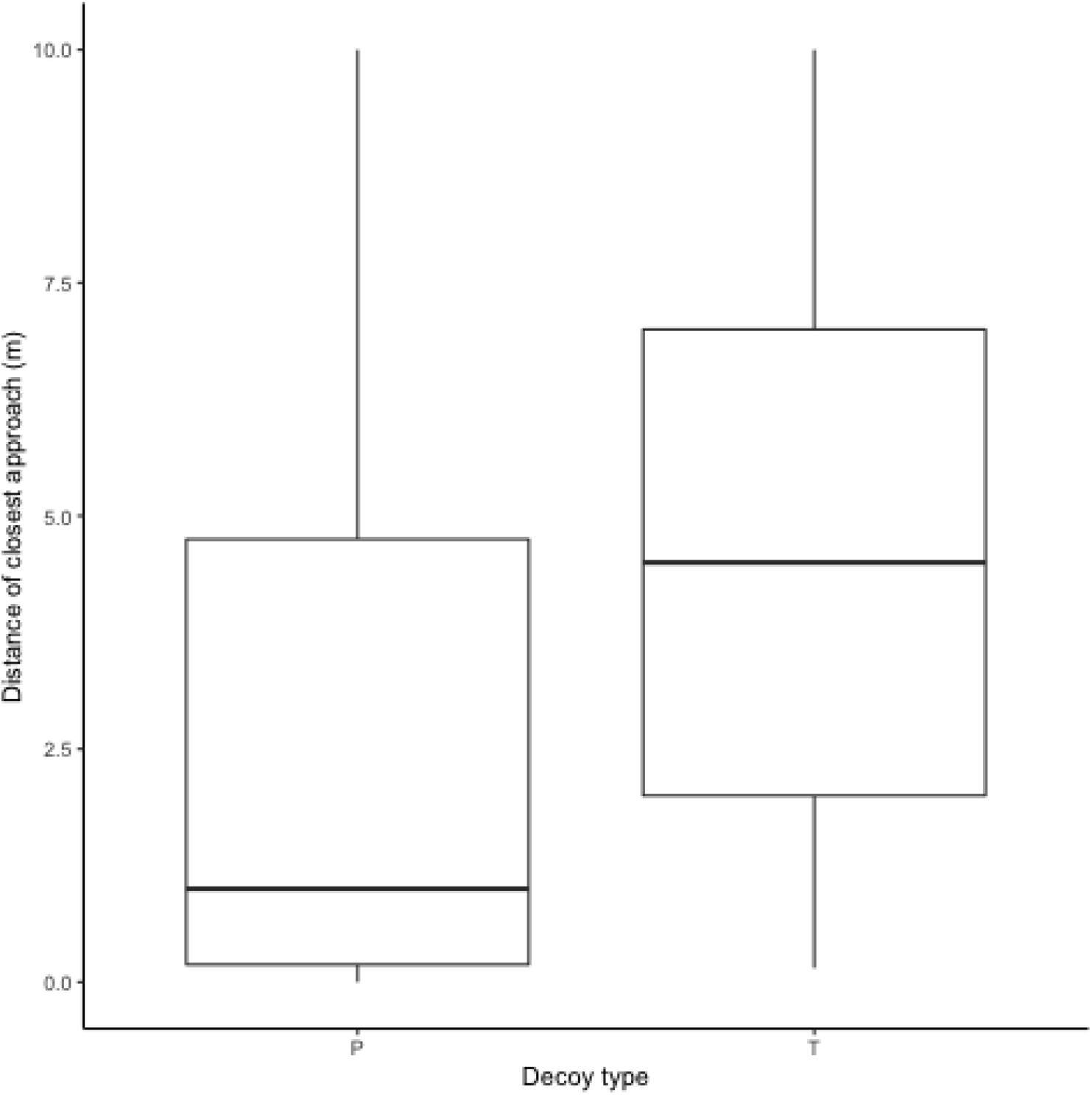
The distance of closest approach of birds on 3D-printed (P) and taxidermied (T) decoys.

**Figure 3.**
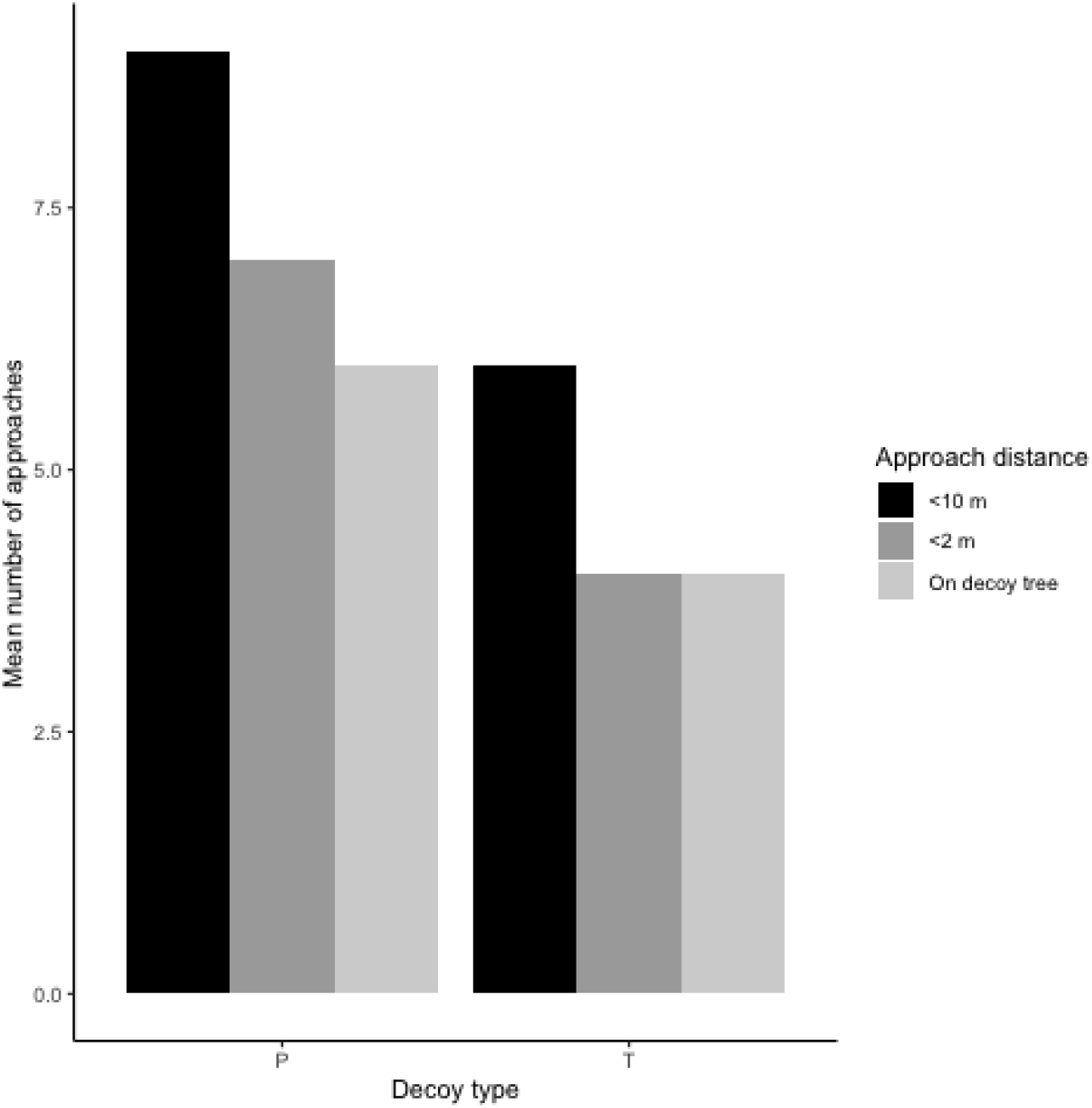
Mean number of approaches of birds within 10 m of the decoy (black), 2 m of the decoy (dark grey), and on the tree containing the decoy (light grey) for the 3D-printed (P) and taxidermied (T) decoys.

## Discussion

Red-breasted sapsuckers are equally likely to approach a plastic 3D-printed decoy as they are a taxidermied conspecific. Our observations suggest they approach closer to a 3D-printed decoy than they do a traditional taxidermied decoy. This pattern is unlikely to be driven by decoy material, rather, we believe our 3D-printed decoy tended to draw sapsuckers nearer as a result of its more lifelike behavioural mimicry. The customizability of 3D design and printing allowed us to imitate woodpecker-specific aggression postures through our decoy, which may have compelled territorial sapsuckers nearer. Relative to nonaggressive live-bird lures, taxidermied decoys posed in permanently aggressive postures induce a stronger territoriality response and higher corticosterone stress levels in affected birds (Scriba & Goymann, 2008). 3D-printed decoys that can imitate aggression have potential to be more effective at attracting birds for capture in mist nets and may either increase the user’s overall number of netted birds, decrease the required time for a successful netting effort, or both.

Behavioural mimicry aside, a sapsucker could simply have perceived the 3D-printed decoy as more lifelike than the taxidermied decoy. Taxidermy is an art, and accurate representation of a specimen’s body shape and posture rely on the skill of the artist. Specimens for a desired species are often in limited supply (Foster & Cannell, 1990), and mistakes in the taxidermy process are permanent. Further, carcasses suffer degrading colouration with age (McNett & Marchetti, 2005; Doucet & Hill, 2009) that could dilute the believability of a taxidermied decoy. In contrast, 3D design facilitates trial and error, and mistakes may be refined without limit; the process does not require artistic ability. A trial and error approach similarly applies to decoy finishing techniques such as painting. Author FT identifies as a poor taxidermist and poorer painter but has produced lifelike 3D-printed decoys for multiple bird species (pers. obs.).

Plastic is a common material in game bird decoys (e.g. Forbes, 1985; Gazalski, 1993), however, such decoys are unlikely to be subjected to the up-close scrutiny of decoys employed in landbird capture. Male song sparrows (*Melospiza melodia*) responded less aggressively to 3D-printed decoys as they did to more-lifelike taxidermied mounts (Beck et al. 2019). FDM-style 3D printing produces visible layer lines (0.2 mm thickness) and may fail to reflect light naturally, reducing the potency in decoy-wildlife interactions (Domingue et al. 2014); however, the lack of significant differences in our decoy treatment suggest FDM-printed plastic has no visual qualities that deter red-breasted sapsuckers. Taxidermied birds pose one significant advantage over hand-painted decoys in that they naturally display plumage coloration in the ultraviolet (UV) region. It is unlikely that UV reflectance plays a major role in red-breasted sapsucker plumage or signalling (Eaton & Lanyon 2003), however, UV coloration may require consideration when imitating some species. Hard decoy materials have the capacity to injure the bills of birds in the course of their provoked aggression (Slevin et al. 2016), though this is of minimal concern for the robust bills of woodpeckers. While the hardness of many solid plastics is comparable to that of the injurious wooden decoys (Slevin et al. 2016), 3D-printed plastic hardness can be functionally reduced through design (Böğrekci et al. 2019). We note that Slevin et al. (2016) acknowledge their publication as the only reported instance of decoy-mediated bill injury, but the risks of decoy hardness may warrant further investigation.

Powerful 3D design software is freely available, and community-driven educational resources abound. Printing services are increasingly common in universities, public libraries, community makerspaces, or by paid private commission. We operated our 3D-printed decoy for more than 16 weeks with no perceivable damage or loss of function. 3D-printed decoys are accessible, affordable, durable, and effective for use in targeted avian mist-netting.

## Declarations

## Acknowledgements

Thank you to Darren Irwin, Ben Freeman, and Rodney Siegel for input on study design. Ildiko Szabo helped with taxidermied decoy construction. Thanks Afnan Ali, Jamie Clarke, and Rashika Ranasinghe for assistance with data collection.

## Author contributions

LN made the stuffed decoy, conducted decoy trials in the field, ran statistical analyses, and contributed to study design and manuscript writing. FT designed and built the 3D-printed decoy and contributed to study design and manuscript writing.

